# ComPath: An ecosystem for exploring, analyzing, and curating mappings across pathway databases

**DOI:** 10.1101/353235

**Authors:** Daniel Domingo-Fernández, Charles Tapley Hoyt, Carlos Bobis-Álvarez, Josep Marín-Llaó, Martin Hofmann-Apitius

## Abstract

Although pathways are widely used for the analysis and representation of biological systems, their lack of clear boundaries, their dispersion across numerous databases, and the lack of interoperability impedes the evaluation of the coverage, agreements, and discrepancies between them. Here, we present ComPath, an ecosystem that supports curation of pathway mappings between databases and fosters the exploration of pathway knowledge through several novel visualizations. We have curated mappings between three of the major pathway databases and present a case study focusing on Parkinson’s disease that illustrates how ComPath can generate new biological insights by identifying pathway modules, clusters, and cross-talks with these mappings. The ComPath source code and resources are available at https://github.com/ComPath and the web application can be accessed at http://compath.scai.fraunhofer.de/.

## Introduction

The notion of pathways enables the representation, formalization, and interpretation of biological events or series of interactions. Cataloging biological knowledge into pathways reduces complexity from all possible interacting molecular entities to a set of well-studied and validated functional relationships between molecular entities culminating in biological processes. Several efforts have generated databases of pathways with varying specificity and granularity that comprise signaling cascades, metabolic routes, and regulatory networks from precise signatures with no more than a couple of acting players to general pathways involving thousands of molecular players (Kanehisa *et al*., 2016; Fabregat *et al*., 2017; Slenter *et al*., 2017; Liberzon *et al*., 2011).

Simplifying biology into pathways and representation as network models or mathematical models inevitably results in a loss of information such as spatiotemporal information or even entire biological entity types. The network abstraction facilitates pathway visualization and interpretation thanks to the harmony between biological networks and systems: nodes correspond to molecular entities and edges to types of interactions occurring between them (e.g., inhibition, phosphorylation, etc.). Although networks can comprise a broad range of molecular types (e.g., proteins, chemicals, small molecules, etc.), they are generally reduced to the most direct outcome of our genetic makeup - the genetic and protein levels - so that we can mechanistically understand their functionality. Thus, they are frequently viewed and simplified to “gene sets”, the collection of all genes/proteins that constitute the pathway, due to the major challenges of incorporating network topology and translating the variety of relationships into pathway analysis methods.

While dedicated research groups and commercial entities with experienced curators have lead a majority of the efforts to compile, delineate, and store biological knowledge into pathway databases (Fabregat *et al*., 2017; Krämer *et al*., 2013), community and crowdsourced efforts have recently gained traction (Slenter *et al*., 2017; Kutmon *et al*., 2016). Further, the variability in curation team composition, database scope (e.g., signaling pathways, gene regulatory networks, and metabolic processes), and curation guidelines led to the adoption of different (and in many ways incompatible) schemata and formalisms such as Biological Pathway Exchange (BioPAX; Demir *et al*., 2010) and Systems Biology Markup Language (SBML; Hucka *et al*., 2003). These incompatibilities motivated the integration and harmonization of resources into pathway meta-databases such as Pathway Commons (Cerami *et al*., 2010) and PathCards (Belinky *et al*., 2015), which focus on integrating databases; iPath (Yamada *et al*., 2011), which focuses on pathway visualization; and SIGNOR, which focuses on signaling pathways (Perfetto *et al*., 2015).

Even after integrating multiple pathway databases into a pathway meta-database, it is difficult to assess the agreements, discrepancies, redundancy, and the complementarity of their contents because of the lack of availability of pathway mappings (e.g., pathway A from resource X is equivalent to pathway B from resource Y) in the original databases. These mappings are difficult to establish because of the arbitrary and overlapping nature of pathway boundaries as well as the absence of a common pathway nomenclature. Several controlled vocabularies have been generated as initial attempts to standardize pathway nomenclature (Petri *et al*., 2014; Iyappan *et al*., 2016), but most pathway databases had already been established by the time these ontologies were published. Therefore, consolidating pathway knowledge is a persisting issue and it is still required to map pathways from different resources together to improve database interoperability.

Hierarchical clustering approaches have been presented as a way of grouping similar pathways based on their corresponding gene sets in order to propose pathway mappings (Belinky *et al*., 2015; Doderer *et al*., 2012). Though these approaches can systematically cluster pathways from multiple resources, there are some limitations to consider: first, the usual tradeoff between over/under-clustering (Daniels and Giraud-Carrier, 2006), and second, pathway nomenclature and biological context are not considered by the clustering algorithm; it often leaves out equivalent pathways with low similarity and ignores the context of the pathway (e.g., cell/disease specificity). Nevertheless, these limitations can be overcome by following clustering and prioritization methods with the manual curation required to interpret the abstract concepts that inherent to pathway definitions (e.g., biological process, cellular location, condition, etc.).

Though numerous algorithms (Khatri *et al*., 2012) and tools (Liberzon *et al*., 2011; Kuleshov *et al*., 2016) have been successfully applied to interpret experimental data through the context of pathway databases (Cary *et al*., 2005; Subramanian *et al*., 2005), there has not yet been a systematic comparison between the contents of various pathway databases, an assessment of their overlaps and gaps, or an establishment of mappings.. Previous studies have only focused on comparing a single or small set of well-established pathways across multiple resources (Bauer-Mehren *et al*., 2009; Chowdhury and Sarkar, 2015). For example, a comparison focused on metabolic pathways revealed how a set of five databases only agreed in a minimum core of the biochemistry knowledge (Stobbe *et al*., 2011).

These studies demonstrate the need to connect insights provided by each pathway database to foster a greater understanding of the underlying biology. Here, we present ComPath, a web application that integrates content from publicly accessible pathway databases, generates comparisons, enables exploration, and facilitates curation of inter-database mappings.

## Results

We developed an interactive web application that enables users to explore, analyze, and curate pathway knowledge. Below, we present three case studies illustrating how it can be used for each of these purposes. The figures for each were generated by interactive, dynamic views in the ComPath web application based on three major public pathway databases: KEGG, Reactome, and WikiPathways **(Figure 1).**

**figure 1.**
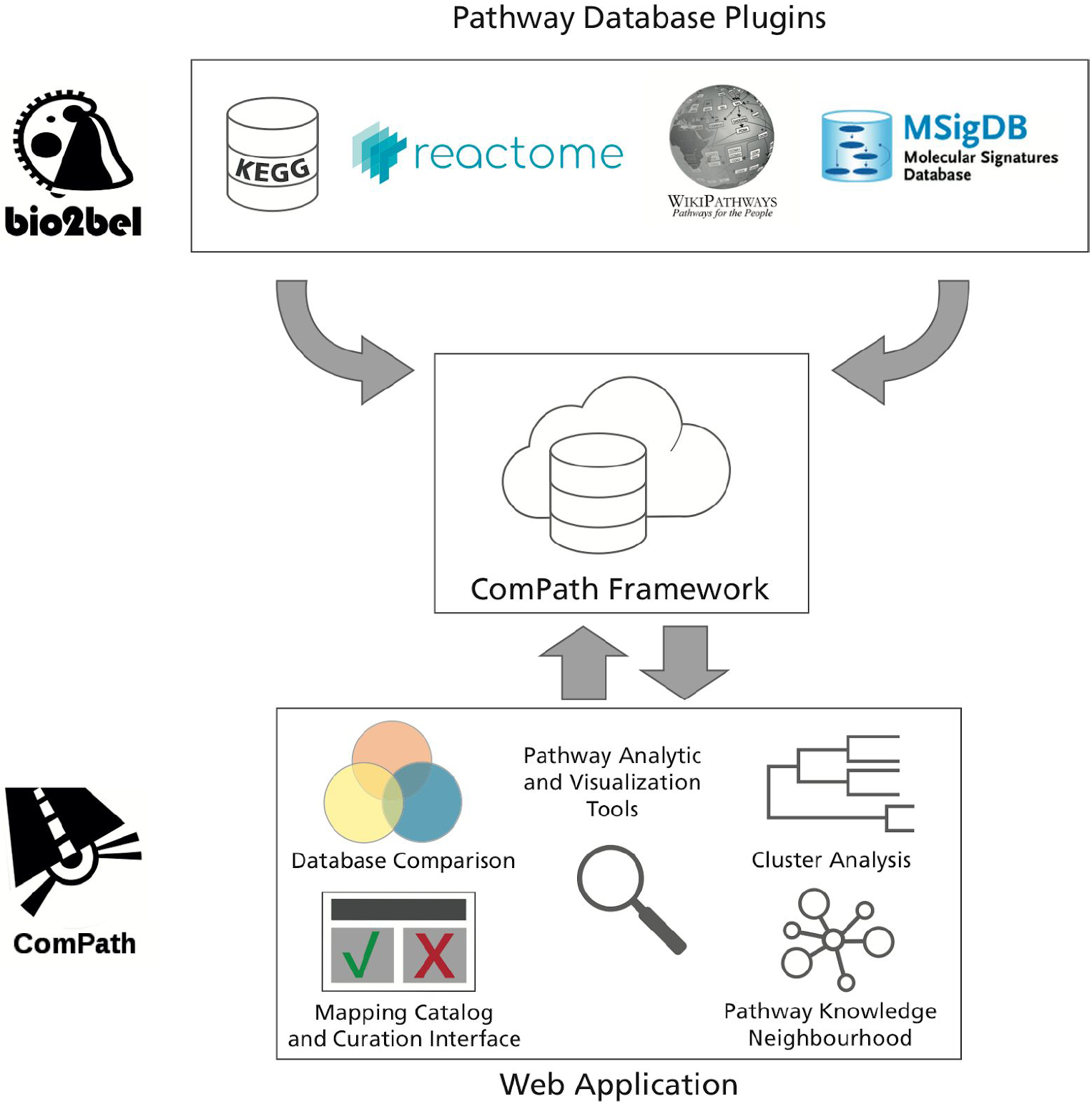
The ComPath ecosystem has three main components: the pathway database plugins, the ComPath framework, and the ComPath web application. The ComPath framework mediates the communication between the plugins containing the pathway database information and the web application.

### Case Study I: Comparison of Pathway Databases

#### Assessment of Gene Coverage

Analysis of the overlaps between Kyoto Encyclopedia of Genes and Genomes (KEGG), Reactome, and WikiPathways revealed that there are approximately 3800 common human genes shared between the three databases (**Figure 2A**). While at least one common human gene was present in almost every pathway across each database, the number of pathways with more common human genes diminishes much more quickly in WikiPathways and Reactome (**Supplementary Figure S1**). This may be due to database properties such as pathway size (e.g., on average, pathways contain 90 genes in KEGG, 50 in Reactome, and 42 in WikiPathways) or gene promiscuity (i.e. genes functionally linked to many pathways) that might influence the results of analyses using pathway resources (**Supplementary Table 2**). For further investigation, the ComPath web application generates summary tables and creates several visualizations to enable exploration of the distributions of pathway size and gene memberships for each database, visualizations that present an overview of the database properties to help identify effects such as gene promiscuity or differences the distribution of gene set sizes (**Figure 2B**).

**figure 2.**
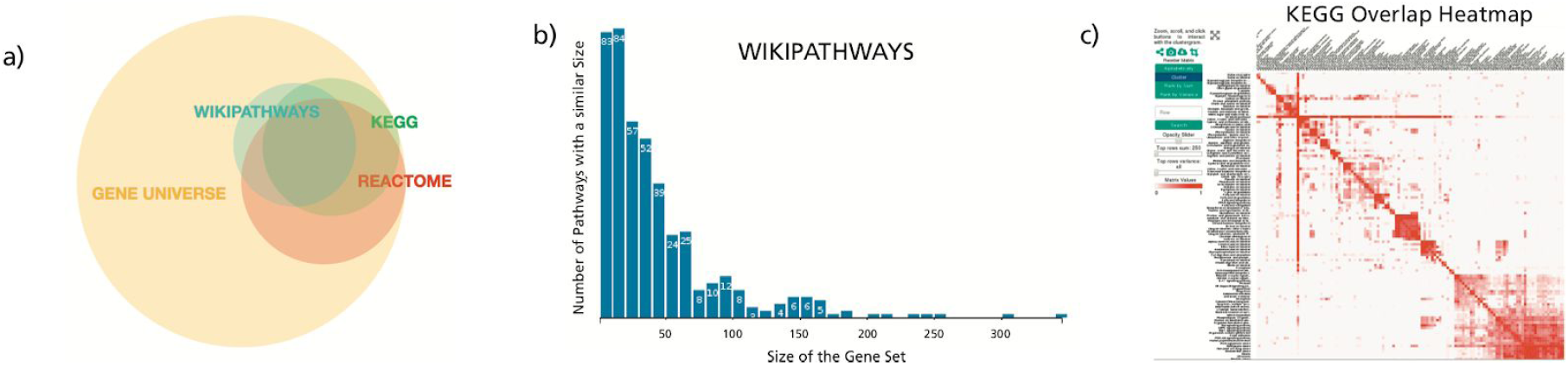
**A)** An Euler diagram summarizing the human gene-centric coverage of KEGG, Reactome, and WikiPathways compared to the universe of all genes from HGNC (more details in **Supplementary Table 1**). **B)** Histogram views present gene promiscuity or pathway size distributions. **C)** The pathway similarity landscape of KEGG visualized as a heatmap.

#### Exploration of Pathways

While the previous views produced gene-centric summaries of the contents of pathway databases, ComPath also enables the exploration of pathway similarity landscape using Clustergrammer.js (Fernandez *et al*., 2017). **Figure 2C** illustrates how this view can identify clusters of pathways based on their similarity and then elucidate the hierarchical relationships between the *Metabolic* pathway, the largest KEGG pathway, and other more high-granular KEGG metabolic pathways (e.g., *alpha-Linolenic acid metabolism, Lipoic acid metabolism, and ether lipid metabolism*).

### Case Study II: Identification of pathway modules, overlaps, and interplays using pathway enrichment

ComPath couples classic pathway enrichment analysis (Kuleshov *et al*., 2016; Reimand *et al*., 2016; Pathan *et al*., 2015; Huang *et al*., 2007) with pathway-centric visualizations to identify modules, investigate overlaps, and cluster pathways. This case study demonstrates their use to investigate the roles of the pathways related to established genetic associations in the context of Parkinson’s disease (PD).

Pathway enrichment with Fisher’s exact test using a gene panel associated with PD reviewed by Brás *et al*. (the gene set will be referenced as PDgset) yielded over 300 pathways containing at least one of the panel’s genes (**Figure 3A**). We discarded pathways with fewer than two genes from PDgset, that were larger than 300 genes, or that were not found to be statistically significant (false discovery rate > 5%) after applying multiple hypothesis testing correction with the Benjamini-Yekutieli method under dependency (Benjamini and Yekutieli, 2001).

**figure 3.**
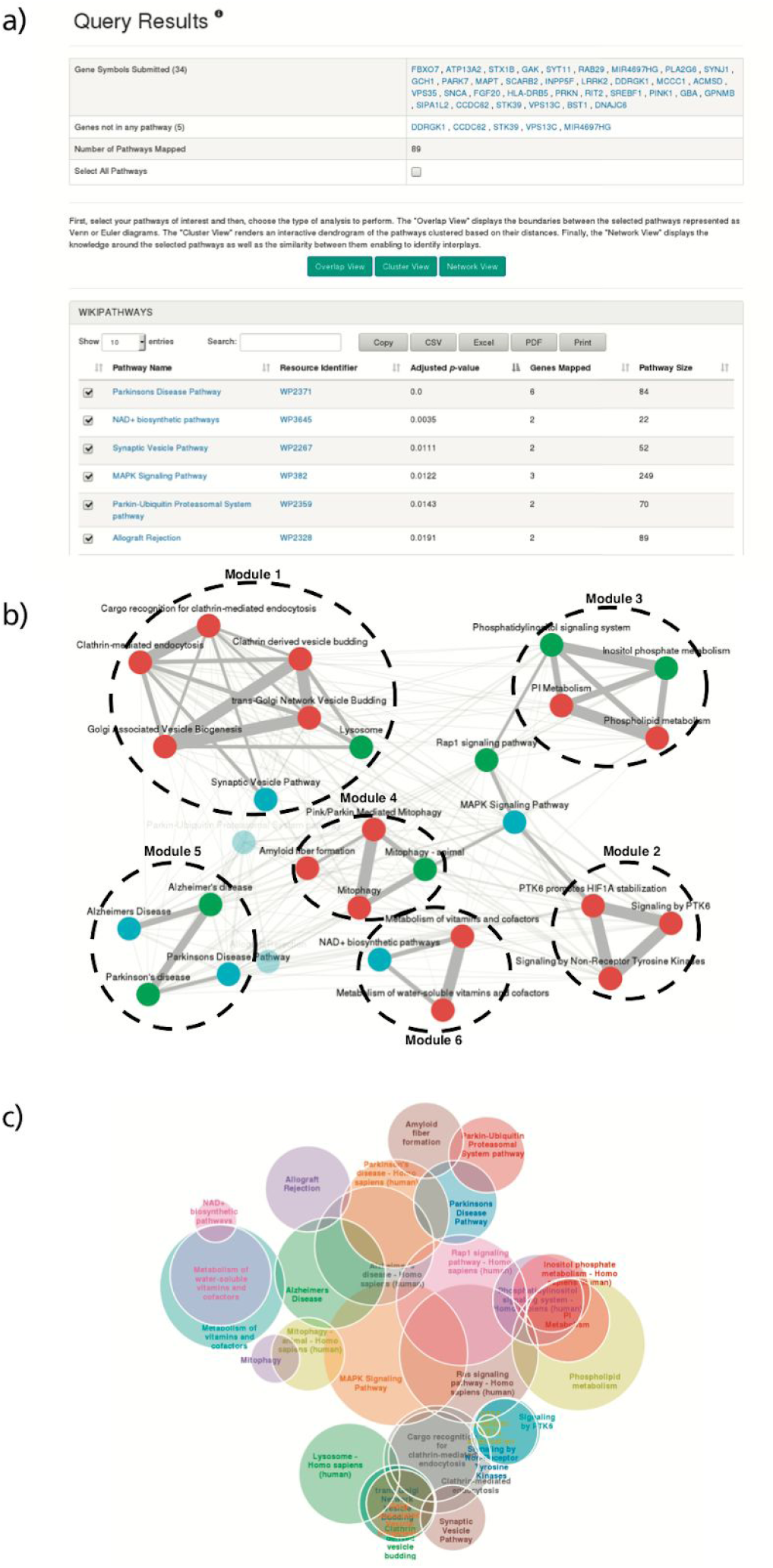
**A)** Results of pathway enrichment using the PDgset as input using the ComPath pathway enrichment wizard. We would like to remark that enrichment results might change over time since ComPath regularly updates their underlying pathway databases. In order to promote reproducibility, the current version of the databases is displayed in the ComPath overview page and older versions can be provided upon request. **B)** The Pathway Network Viewer displays the similarity around a selection of pathways. **C)** The Pathway Overlap View depicts the overlaps and intersection of pathways enriched from the PDgset.

Three views were used to assist in the interpretation of the remaining 29 enriched pathways: a pathway network view was used to identify pathway modules, a pathway overlap view was used to explore the intersections and cross-talks between pathways, and a pathway dendrogram view was used for clustering.

The pathway network view renders a pathway-to-pathway network in which nodes represent pathways and weighted edges represent their corresponding gene set similarities in a similar fashion to PathwayConnector (Minadakis *et al*., 2018). For the PDgset, this visualization helped us to define six different modules (i.e., groups of pathways) by removing edges with a weight lower than 0.2 (**Figure 3B**). The largest module (labelled as M_1_) contained pathways related to the processes of endocytosis and vesicle transport, both of which are putatively disrupted in PD (Perrett *et al*., 2015). M_2_ comprised pathways related to PTK6 signaling such as the Reactome pathway, *PTK6 promotes HIF1A stabilization,* whose high pathway enrichment significance (q-value=0.0005), as well as its role in regulating another PDgset gene, ATP13A2 (Rajagopalan *et al*., 2016), suggests that it may be linked to PD. ATP13A2 is directly responsible for Kufor-Rakeb syndrome (Gusdon *et al*., 2012), a rare juvenile form of PD, and participates in two other PD mechanisms: lysosomal iron storage and mitochondrial stress. Because pathways related to these two mechanisms (i.e., *Lysosome pathway* from KEGG, *Pink/Parkin mediated mitophagy* from Reactome, and *Mitophagy pathway* from both KEGG and Reactome; M4) were also enriched by pathway enrichment analysis, we investigated the role of ATP13A2 in PD further.

ATP13A2isactivatedbyphosphatidylinositol(3,5)bisphosphate, aparticular phosphatidylinositol involved in M_3_ pathways (phosphatidylinositol metabolism and signaling pathways). Because this activation leads to a reduction in mitochondrial stress and α-synuclein toxicity, two hallmarks of PD, ATP13A2 has been proposed as a therapeutic target (Holemans *et al*., 2015). Ultimately, the exploration of the similarities and cross-talks between these three modules suggests further investigation of the candidate PD gene ATP13A2. Ultimately, this view complements pathway enrichment in the identification of pathway modules, exploration pathway cross-talks, and prioritization of genes for further study.

While the pathway network viewer provides an overview of the different modules and their cross-talks, it does not reveal information about their contained pathways’ boundaries and intersections. Therefore, we implemented the pathway overlap view; an interactive Euler diagram that allows exploration of pathway demarcations (**Figure 3C**). We employed this view to identify the set of genes common to all pathways in M_5_, a module comprising the two Alzheimer’s disease (AD) and two PD pathways from KEGG and WikiPathways. Subsequently, we used the ComPath pathway enrichment wizard to investigate in which pathways the common five genes identified (APAF1, CASP3, CASP9, CYCS, and SNCA) participate. The analysis revealed that they are predominantly involved in apoptosis, an important process in both AD and PD pathophysiology (Obulesu *et al*., 2014; Tatton *et al*., 2003).

The third visualization renders the results of the hierarchical clustering approach described in Chen *et al*. in the form of a dendrogram, enabling deterministic pathway grouping based on gene set similarity. We used this view in the PDgset example to assign the pathways without module membership to the closest module (**Supplementary Figure S2**). The dendrogram proposed merging three previously unassigned pathways into M_2_ (i.e., *Allograft Rejection, MAPK Signaling pathway,* and *Rasp1 signaling pathway*). Additionally, the resulting dendrogram from clustering revealed hierarchical relationships between pathways (e.g., *Pink/Parkin Mediated Mitophagy* is a subset of the Reactome *Mitophagy* pathway), information that can be used to establish pathway mappings, as we show in the following case study.

#### Case Study III: Establishing mappings between pathway databases

ComPath, as well as other tools, have demonstrated the benefits of integrating pathway knowledge from diverse resources to improve biological functional analysis (Cerami *et al*., 2010; Belinky *et al*., 2015; Kuleshov *et al.,* 2016). However, even after overcoming the technical hurdle of harmonizing different formats used by different databases, these integrative approaches must be complemented by mappings at a pathway level in order to have cross references between databases; thus, improving their interoperability. Such information could then be used to first link related pathways and then investigate their interplays, explore the consistency of their boundaries, calculate their discrepancies and agreements, or simply contextualize the knowledge around a certain biological process.

In order to address this, ComPath introduces a curation environment in which users from the scientific community can propose and maintain a collection of established mappings between pathways from various databases. This laborious task is facilitated by the interactive visualizations (i.e., a dendrogram view and a similarity landscape heatmap) presented in the previous case studies as well as dedicated pathway pages where the content, descriptions, references, and the established mappings can be examined (**Figure 4A**). Furthermore, ComPath suggests the most similar pathways based on this information so users can propose new mappings. This new mappings are included into the mapping catalog that serves as a search interface as well as a distribution platform for mappings (**Figure 4B**). In addition, the mapping catalog promotes community engaging incorporating a voting system where authenticated users can agree or disagree on mappings; this way, proposed mappings with a net sum of votes greater than 3 are automatically registered as accepted.

**figure 4.**
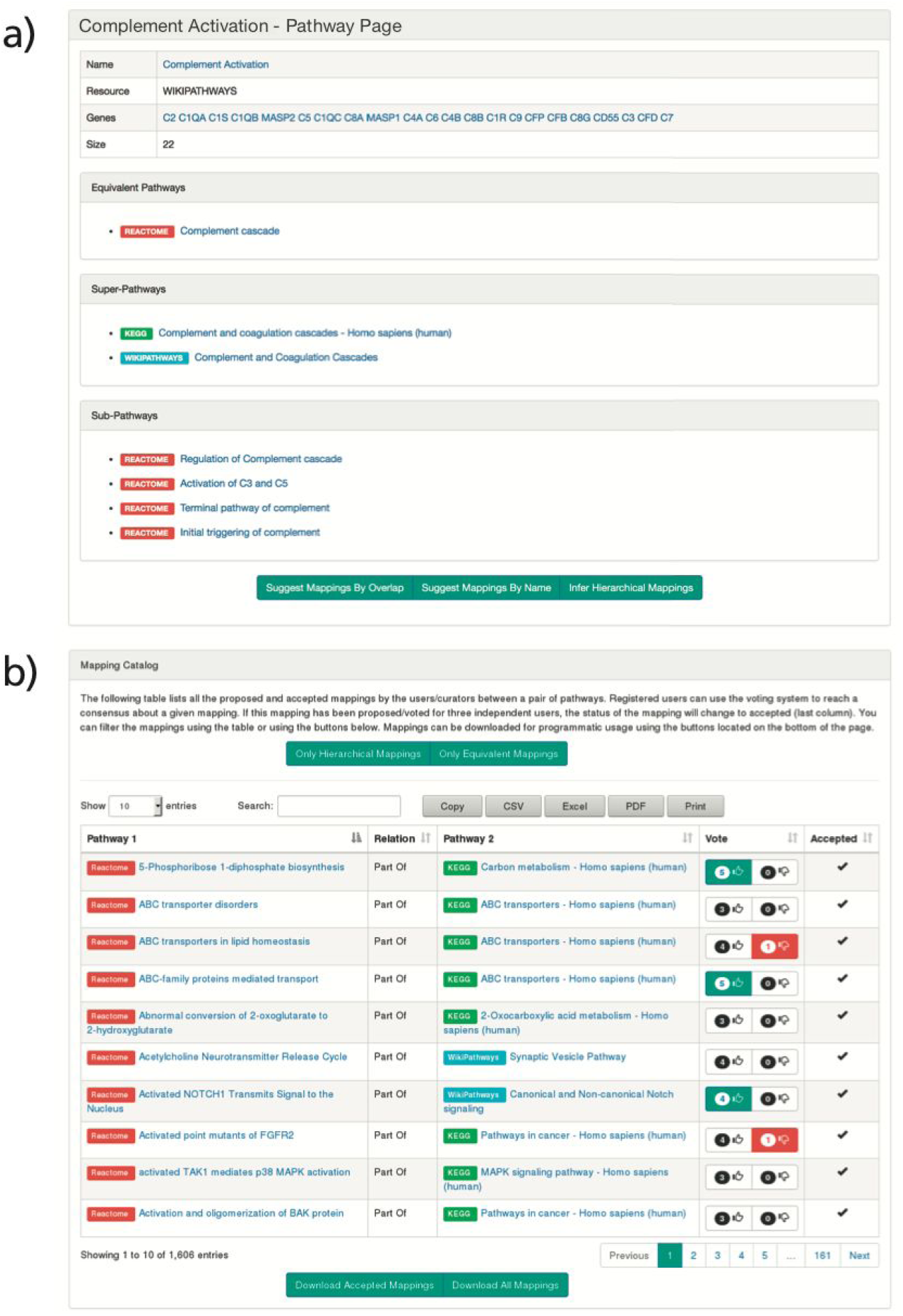
**A)** The pathway info view introduces basic pathway information such as its participating molecular entities, references, or mappings and enables automatic mapping suggestions based on different similarity metrics.

Furthermore, the mappings of the selected pathway can be visualized with a dynamic view that enables exploration of multiple levels of its hierarchy (**Supplementary Figure S3**). **B)** The mappings view allows users to browse established mappings, propose new mappings, and give feedback on putative mappings.

After an exhaustive investigation of all possible mappings between pathways in KEGG, Reactome, and WikiPathways (see Methods), we identified 58 equivalencies between KEGG and Reactome, 64 between Reactome and WikiPathways, and 55 between KEGG and WikiPathways. Of these equivalent pathways, 21 are shared between the three resources (**Figure 5 and Supplementary Table 4**). We also identified 247 hierarchical relationships between KEGG and Reactome, 597 between KEGG and WikiPathways, and 564 between Reactome and WikiPathways. After considering these, approximately 26% of KEGG, 70% of Reactome, and 35% of WikiPathways did not share any mappings with any other database (**Supplementary Figure S4**). The high uniqueness observed in Reactome could be attributed to several factors: its small pathway sizes, its high granularity, and its high coverage of HGNC (**Figure 2A**).

**figure 5.**
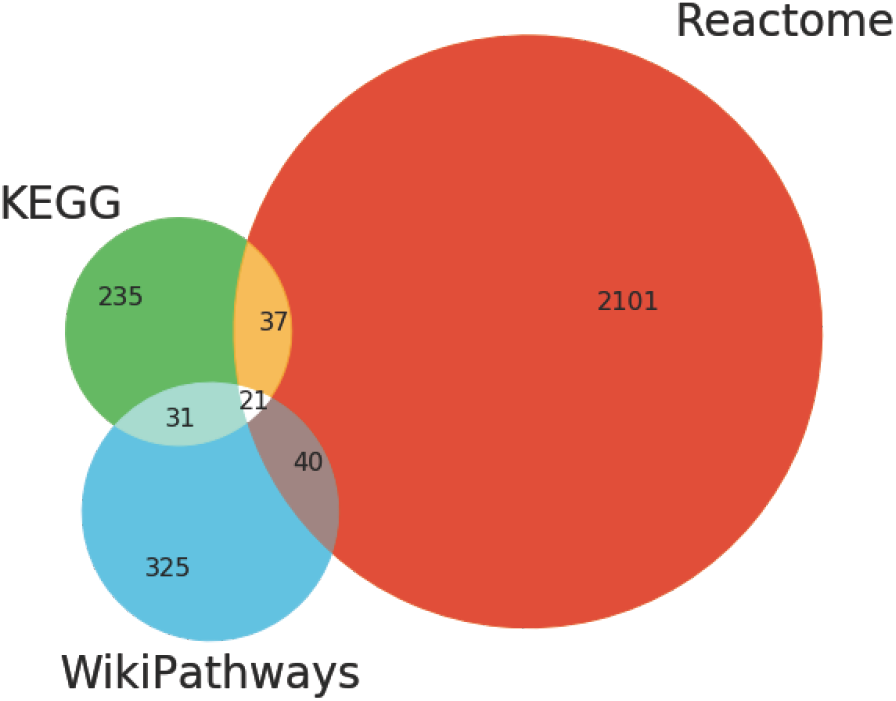
Venn diagram illustrating the overlaps of equivalent pathways between KEGG, Reactome, WikiPathways resulting from the curation exercise. Note: the number of overlapping pathways in the Venn diagram do not exactly match the number of equivalent mappings since there are equivalent pathways within WikiPathways that, when mapped to another database, could have more than one equivalent pathway. For example, there are two equivalent *Wnt signaling pathways* in WikiPathways that are both mapped to their corresponding Reactome pathway. This is resolved to a unique in the Venn diagram. A list of intra-database equivalent pathways is presented in the **Supplementary Table 3.**

The results of this curation effort are distributed at https://github.com/ComPath/resources and https://compath.scai.fraunhofer.de/ so they can be revised, updated, and exploited by the research community hoping that this work serves as a first endeavor towards unifying pathway knowledge.

## Discussion

The lack of a lingua franca in systems biology hampers the harmonization that would enable the exploration of the coverage, agreements, or discrepancies in the pathway knowledge. Harmonizing this information is an important step to better comprehend and model biology as well as improve the bioinformatics pipelines that utilize this knowledge to elucidate biological insights. As a first step towards closing this gap, we have implemented an environment capable of accommodating the pathway knowledge from multiple databases in order to facilitate its exploration and analysis through a web application. The flexibility of ComPath enables the incorporation of additional databases as well as dynamic update of its resources; the latter of which is often neglected, but can have a significant effect on derived analyses (Wadi *et al*., 2016). Additionally, an embedded curation interface allow users to curate and establish mappings between pathways. Accordingly, we used ComPath to conduct extensive curation work to link the pathways from three major pathway databases in order to evaluate their similarities and differences. This mapping catalog serves as a first effort towards unifying and linking pathway information across databases that can later be adopted by the original databases or to create ontologies that store these mappings. Finally, we plan to update the source database biannually as well as curating mappings for newly added pathways.

The common genes between KEGG, Reactome, and WikiPathways covered the majority of pathways, indicating that their pathway knowledge is partially biased towards this shared gene set, even while there are still thousands of genes that have not yet been functionally annotated to pathways. Furthermore, our curation effort revealed that a surprisingly low number of pathways (21) were equivalent between KEGG, Reactome, and WikiPathways. On the other hand, the number of mapped pathways increased significantly when the hierarchical mappings were considered, revealing the inconsistent granularity employed to delineate pathway boundaries.

Although the absence of topological pathway information in ComPath is an irrefutable limitation in this study, gene-centric approaches enable a reduction of complexity in pathway comparison as well as integration of resources which do not provide topology information (Belinky *et al*., 2015). Furthermore, recent studies revealed significant differences across a large sample of topology-based pathway analysis methods (Ihnatova, *et al*., 2018), and highlighted that gene sets alone might be sufficient to detect an enriched pathway under realistic circumstances (Bayerlová *et al*., 2015). Hence, even if the abstraction of pathways as gene sets might not exploit all the existing pathway information, it is sufficient to drive an investigation of the pathway knowledge.

The established inter-database mappings allowed to link pathways from three major databases, opening the door towards a better integration of the pathway knowledge. In the future, these links can be used to complement and fill pathway knowledge as well as to conduct a precise evaluation of equivalent or related pathways by exploiting the available format converters such as the converter from Reactome to WikiPathways (Bohler *et al*., 2016). Furthermore, ComPath have been designed to accommodate multiple types of molecular entities participating in pathways (i.e. Reactome chemical information); thus, enabling to replicate the analyses presented with lipid or metabolite databases such as LIPEA (Acevedo *et al*., 2018) or HMDB (Wishart *et al*., 2017).

In summary, we demonstrated that ComPath serves as an exploratory, analytic, and curation framework for pathway databases. Furthermore, we showed how the ComPath web application can complement enrichment approaches to elucidate and prioritize pathways and genes related to interesting biological phenomenon. Finally, we hope that the implementation of a curation ecosystem and the first mapping efforts conducted in this work pave the way towards unifying the pathway knowledge.

## Methods

### Implementation

#### ComPath Framework

At its core, ComPath is a framework for integrating pathway and gene set databases. We defined a set of guidelines for implementing wrappers around the processes of downloading data, transforming it into a common data model, and making queries. These guidelines are encoded in an abstract class with the Python programming language such that new plugins can be quickly implemented for new resources. Each implementation must have a mapping between genes and pathways as well as functions for exporting pathways as gene sets, performing pathway enrichment analysis, and performing reasoning/inference over pathway hierarchies.

#### Compath Plugins

We implemented plugins for four major public pathway databases: KEGG, Reactome, WikiPathways, and MSigDB (Kanehisa *et al*., 2016; Fabregat *et al*., 2017; Slenter *et al*., 2017; Liberzon *et al*., 2011). They can be used individually as a way of extracting, updating, and exploring the pathways contained within the database. Additionally, they can be used jointly in the ComPath web application where the pathways from multiple databases are integrated for their exploration, analysis, and curation.

#### ComPath Web Application

The web application was implemented in the Python programming language using the Flask microframework and a suite of its extensions. The compatibility between Flask and the data models defined in all pathway plugins allows the integration and harmonization of the pathway knowledge in an extensible manner. To illustrate the flexibility of ComPath, we have included plugins for the Alzheimer’s disease and Parkinson’s disease gene sets associated with disease-specific mechanisms from NeuroMMSig (Domingo-Fernández *et al*., 2017) in the public version of the ComPath web (https://compath.scai.fraunhofer.de/).

ComPath leverages a variety of state-of-the-art libraries for visualization and exploration of pathway knowledge. We chose Bootstrap for the design of the website since its responsive design retains full compatibility across all devices. Interactive visualizations are generated using several Javascript libraries, including D3.js, Clustergrammer.js (Fernandez *et al*., 2017), and Cytoscape.js (Franz *et al*., 2015).

We implemented a RESTful API documented with an OpenAPI specification that can be accessed through the ComPath instance released at https://compath.scai.fraunhofer.de/apidocs. The API enables users to programmatically extract mapping information and perform queries using different genes or pathways identifiers.

#### Code Availability

The source code for ComPath and its plugins can be found on GitHub (https://github.com/ComPath and https://github.com/Bio2BEL) under the MIT license. Both the plugins and the web application can be installed with PyPI (https://pypi.org), the main packaging system for Python. Furthermore, we have included a Dockerfile to enable reproducing the ComPath environment with Docker (https://www.docker.com/). Finally, documentation is included in each GitHub repository and it is also accessible at Read the Docs (https://readthedocs.org).

## Methods

### Estimating Pathway Similarity

While a variety of indices (e.g., Jaccard, Sørensen–Dice, Tversky) have been used to assess the similarity between sets, the Szymkiewicz-Simpson coefficient (Equation 1) is most appropriate for comparing sets widely varying in size. Similarly to previous studies, we have chosen this index to not only calculate pathway similarity but also reveal *contained* pathways (i.e., when most of the nodes from a small pathway are in a larger pathway) to indicate potential hierarchical relationships (Chen *et al.,* 2014, Pita-Juarez *et al*., 2018; Belinky *et al*., 2015; Katiyar *et al*., 2018)

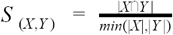

Equation 1. The Szymkiewicz-Simpson coefficient calculates the similarity between two sets (X and Y) where 0 ≤ S ≤ 1. The similarity is the size of the intersection of the two sets divided by the size of the smaller.

### Curation of Pathway Mappings

Here, we describe the curation procedure we used in order to systematically generate equivalency and hierarchical mappings between the human pathways originating from KEGG, Reactome, and WikiPathways. Here, it is important to note that we have only focused on generating mappings for the pathways originating from each of the three resources, not their imported pathways from other databases (e.g., WikiPathways imported Reactome pathways that are evidently equivalent to the ones in Reactome). First, we define two types of mappings:

1. **equivalentTo**. An undirected relationship denoting both pathways refer to the same biological process. The requirements for this relationship are:
  - *Scope*: both pathways represent the same biological pathway information.
  - *Similarity*: both pathways must share at minimum of one overlapping gene.
  - *Context*: both pathways should take place in the same context (e.g., cell line, physiology).
  - **isPartOf**. A directed relationship denoting the hierarchical relationship between the pathway 1 (child) and 2 (parent). The requirements are:
    - *Subset Scope*: The subject (pathway 1) is a subset of pathway 2 (e.g., Reactome pathway hierarchy).
    - *Similarity*: same as above.
    - *Context:* same as above.

We generated all possible mappings between pathways in each database (KEGG-WikiPathways, KEGG-Reactome, and WikiPathways-Reactome) and prioritized them based on the follow two independent metrics that have been proposed to calculate pathway similarity (Belinky *et al*., 2015):

1. Lexical similarity between each pair of pathways’ names was calculated using the Levenshtein distance (Levenshtein, 1966).
2. Content similarity between each pair of pathways’ genes was calculated using the previously described Szymkiewicz-Simpson coefficient.

After prioritization, our three curators from different areas of expertise (neuroscience, medicine, and biology) independently evaluated both similarities and the scope and context included in the pathway descriptions to assign the mapping types and to remove false positives. Furthermore, we investigated possible intra-database mappings within KEGG and WikiPathways since these resources do not yet contain hierarchical relationships. Finally, our curators combined the results and re-evaluated them to generate a consensus mapping file. It is available at https://github.com/ComPath/resources under the MIT License.

## Acknowledgements

This work was supported by the EU/EFPIA Innovative Medicines Initiative Joint Undertaking under AETIONOMY [grant number 115568], resources of which are composed of financial contribution from the European Union’s Seventh Framework Programme (FP7/2007-2013) and EFPIA companies in kind contribution.

## Author Contributions

M.H.A and D.D.F conceived and designed the study. D.D.F implemented ComPath and the pathway database plugins with help from C.T.H. D.D.F, C.B.A, and J.M.L curated the pathway mappings. D.D.F and C.T.H wrote the paper. M.H.A. reviewed the content.

